# Live detection of neural stem and glioblastoma cells by an oligothiophene derivative

**DOI:** 10.1101/579185

**Authors:** Shirin Ilkhanizadeh, Aileen Gracias, Andreas K.O. Åslund, Marcus Bäck, Rozalyn Simon, Vilma Rraklli, Bianca Migliori, Edel Kavanagh, Sven Nelander, Bengt Westermark, Lene Uhrbom, Karin Forsberg-Nilsson, Ana I. Teixeira, Peter Konradsson, Per Uhlén, Bertrand Joseph, Ola Hermanson, K. Peter R. Nilsson

## Abstract

Here we report a luminescent conjugated oligothiophene (LCO), named p-HTMI, for non-invasive and non-amplified real-time detection of live human patient-derived glioblastoma (GBM) cells and embryonic neural stem cells (NSCs). While p-HTMI stained only a small fraction of other cell types investigated, the mere addition of p-HTMI to the cell culture resulted in efficient detection of NSCs or GBM cells from rodents and humans within minutes. p-HTMI is functionalized with a methylated imidazole moiety resembling the side chain of histidine/histamine, and non-methylated analogues were not functional. p-HTMI were readily able to detect a subpopulation of human cells *in vivo* in mouse brain sections with tumors developed from orthotopically transplanted patient-derived GBM cells. Cell sorting experiments of human GBM cells demonstrated that p-HTMI labeled the same population as CD271, a proposed marker for stem cell-like cells and rapidly migrating cells in glioblastoma. Our results suggest that the LCO p-HTMI is a versatile tool for immediate and selective detection of subpopulations of neural stem and glioma cells.

---

Small probes specific for a distinct biomolecule or structural element have proven useful for recording biological events. In this regard, luminescent conjugated oligo- and polythiophenes (LCOs and LCPs) have been utilized as target-specific chameleons that change their emission depending on the structural motif of a distinct target molecule even in a complex environment such as tissue [1]. LCOs and LCPs contain a repetitive flexible thiophene backbone, and a conformational restriction of the thiophene rings upon interaction with a biomolecule leads to a distinct optical fingerprint from the dyes. Previously, chemically defined anionic pentameric LCOs have been reported as amyloid specific ligands for optical detection of pathogenic protein aggregates *in vivo* [2].

Stem cells derived from the developing and adult brain (neural stem cells; NSCs) and cell populations derived from glioblastoma (GBM) tumors have caught special attention due to the need for new approaches in diagnosis, treatment, and possible cures for diseases of the nervous system [3]. GBM is an aggressive type of nervous system tumor with a mean survival time of 12-14 months, and the survival time has remained low in spite of major efforts [4]. At least one subpopulation of progenitors with stem cell-like properties can be derived from GBM tumors, and these cells are referred commonly to as glioblastoma-derived stem cell-like cells (GSCs) [5]. GSCs seem to escape conventional irradiation treatment, chemotherapy, and surgery [4], and it is therefore urgent to develop novel approaches for reliable detection of these and other cell types in GBM. While genetic approaches have proven successful in identifying and deciphering progenitors and stem cells in the healthy nervous system [6], simple and reproducible techniques to rapidly identify and detect live NSCs and GBM cells without invasive or genetic modulation are desirable [7]. Most commonly, either antibodies or tools like green fluorescent protein (GFP) are used for detection of specific stem cell types, but these modes have several limitations for detection of live cells as they often are time consuming, require secondary amplification, and/or manipulation of the cells and/or organism.

Herein, we investigate the use of a library of LCOs with distinct side-chain functionalization as molecular probes for specific identification and detection of NSCs and GBM-derived cells in real time.

In order to characterize the library of LCOs in a NSC context, we used several very well-established differentiation protocols to generate a variety of cell types from NSCs derived from embryonic rat cortices to compare with the undifferentiated cells. This NSC protocol, based on fibroblast growth factor 2 (FGF2) treatment, is well-characterized and widely used [8] and has been shown to contain of ≫99% cells with ability to differentiate into neurons, astrocytes, and oligodendrocytes [8a,9]. Interleukin-6-related cytokines such as the extrinsic factor ciliary neurotrophic factor (CNTF) induces astrocytic differentiation, fetal bovine serum (FBS) induces mesenchymal differentiation (smooth muscle cell-like), and a combination of BMP4/Wnt3a induces neuronal and astrocytic differentiation [10]. In a screen with various LCO libraries, we identified p-HTMI (**Scheme 1, Figure 1a, Supplementary Figure S1**) as a potential marker for NSCs. Within 10 minutes after application of p-HTMI by simply pipetting into the cell culture medium, fluorescence at a wavelength common to green fluorescent proteins was generated in undifferentiated embryonic rat NSCs (**Figure 1b**). Two-photon microscopy revealed a strong fluorescent signal accumulated in the cytoplasm of the cells, thus indicating the probe had efficiently crossed the cell membrane (**Figure 1d, Supplementary Movie 1**). Simultaneous detection of a chloromethyl derivative detecting all live cells with red fluorescence (CellTracker) demonstrated that p-HTMI-stained cells were viable. The red and green fluorescence could be detected in parallel in the same cells, and thus p-HTMI allowed double-labeling of single cells (data not shown, see further below). The differentiated and more mature cells displayed a significantly lower or no signal (**Figure 1b**), and testing of a long series of various cells, including several commonly used cell lines (HEK-293, CV-1, COS-7 etc), mesenchymal stem cells, or embryonic stem cells (see further below) did not result in any staining by p-HTMI. In contrast, the variant LCO p-HTE-Ser (**Supplementary Figure S1**) displayed no signal in undifferentiated NSCs whereas a weak staining was observed in fully differentiated smooth muscle cells and mature astrocytes (**Supplementary Figure S2**). Anionic LCPs and LCOs (PTAA, p-FTAA, p-HTAA, h-HTAA (**Supplementary Figure S1**)) previously reported to specifically detect amyloid structures [6b,11], displayed no cell-specific signal in undifferentiated NSCs (data not shown).

As p-HTMI is functionalized with methylated imidazole moieties (**Scheme 1 and Figure 1a**) resembling the side chain of histidine/histamine, we asked whether the specificity of the NSC staining was dependent on this distinct side chain functionalization. We therefore generated pentameric LCOs with imidazole moieties lacking the methylation, p-HTA-His and p-HTIm (**Supplementary Figure S1, see also Scheme 1 and Supplementary scheme 2**). Interestingly, we found that these molecules did not detect the NSCs tested above (**Figure 1c and data not shown**), suggesting that the methylation of the imidazole moieties is a crucial factor regulating the efficiency and specificity of the detection. However, a previously reported polydispersed LCP with methylated imidazole moieties attached to all thiophene units, PTMI [12] (**Supplementary Figure S1**) did not stain the NSCs either, suggesting that the length of the thiophene backbone or the positioning of the methylated imidazole moieties are additional important parameters to obtain the selective staining of the NSCs.

**Figure 1.**
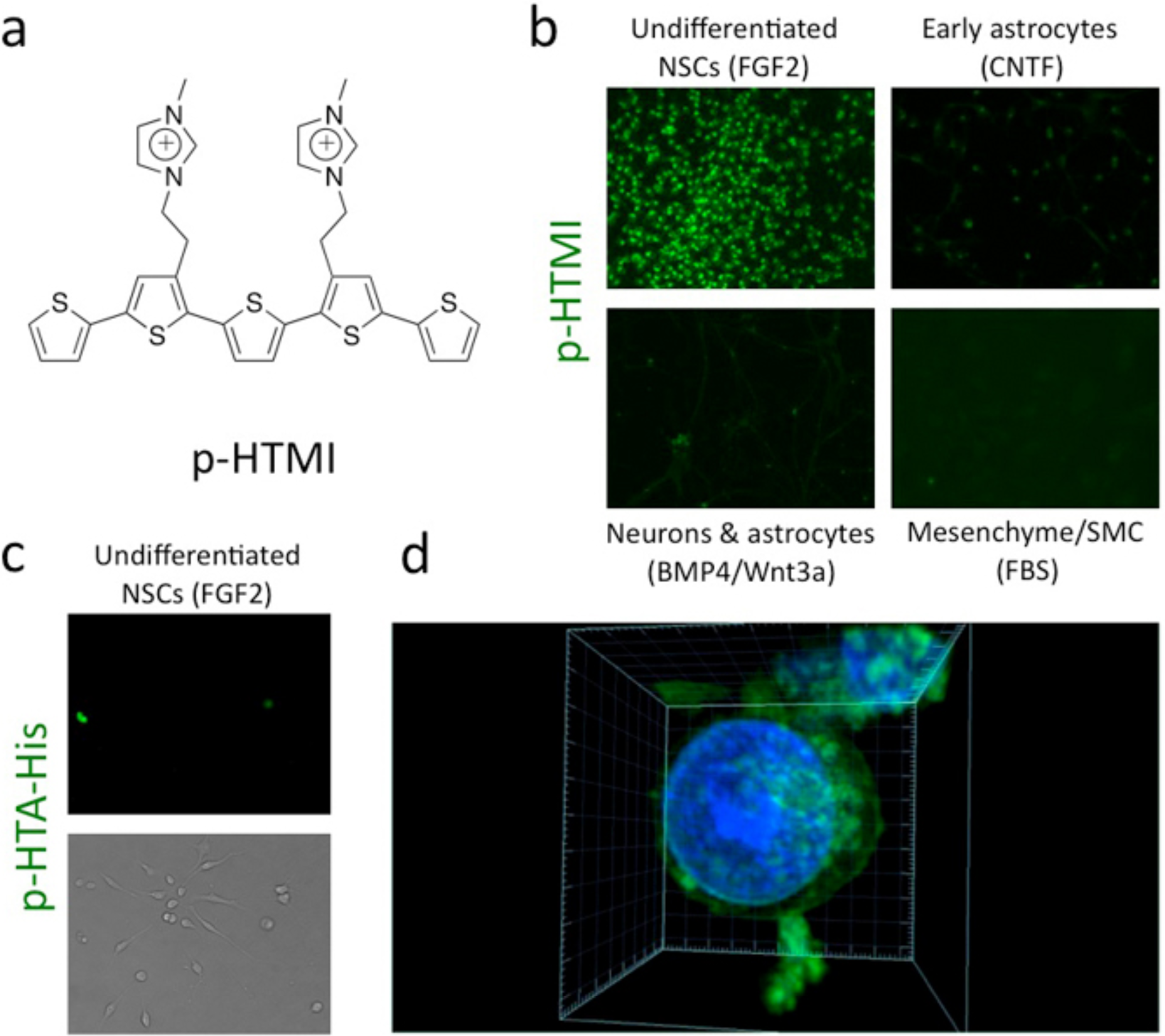
p-HTMI is a novel oligothiophene that specifically detects embryonic neural stem cells (NSCs) *in vitro*. (**a**) The chemical structure of p-HTMI displays a methylated imidazole moiety. (**b**) Micrographs depicting staining of embryonic neural stem cells (NSCs) and cells differentiated thereof. Staining obtained c:a 10 min after administration of p-HTMI (1:500). The staining showed a high signal-to-noise ratio and was predominantly cytoplasmic. (**c**) Micrograph depicting undifferentiated NSCs 10 min after administration of p-HTA-His, an LCO with imidazole moieties lacking the methylation. (**d**) Two-photon microscopy confirmed that the p-HTMI labeling (green) was predominantly cytoplasmic. DAPI in blue labels the cell nucleus DNA. See also Supplementary Movie 1.

Fluorescence activated cell sorting (FACS) is commonly used to quantitatively record fluorescent signals from individual cells and to separate cells based on fluorescence. We stained FGF2-treated undifferentiated NSCs with either p-HTMI or p-HTE-Ser. Using FACS, we then sorted unstained cells, p-HTMI stained cells and p-HTE-Ser stained cells. The cells stained with p-HTE-Ser displayed fluorescence comparable to the unstained cells and this staining was thus considered background fluorescence (**Supplementary Figure S3**). p-HTMI stained cells however displayed a clear fluorescence peak, not overlapping with background fluorescence levels (**Supplementary Figure S3)**. In order to check for cell-to-cell leakage of the molecules, cells separately stained with p-HTMI and p-HTE-Ser were mixed and sorted with FACS. Two distinct fluorescence peaks were obtained, showing that cross contamination of the molecules does not occur (**Supplementary Figure S3**). Taken together, these results verify that p-HTMI selectively detects undifferentiated embryonic NSCs *in vitro*.

We next wanted to test whether the observed p-HTMI staining was a result of the treatment of the cells rather than the NSC phenotype. We therefore used mouse embryonic stem cells (ESCs) or NSCs derived from these cells by using an established retinoic acid-based protocol clearly distinct from the NSC-expanding protocol used hitherto [13]. Intriguingly, while p-HTMI did only stain a small fraction of ESCs (**Figure 2a**), we found staining similar to that of embryonic brain-derived NSCs in the ESC-derived NSCs at similar efficiency (**Figure 2b**). This result indicated that p-HTMI is labeling embryonic or embryonic-like NSCs and not cells cultured with a particular protocol. In addition, p-HTE-Ser labeled >90% of the ES cells but did not show any staining in the ESC-derived NSCs (**Figure 2c and data not shown**).

**Figure 2.**
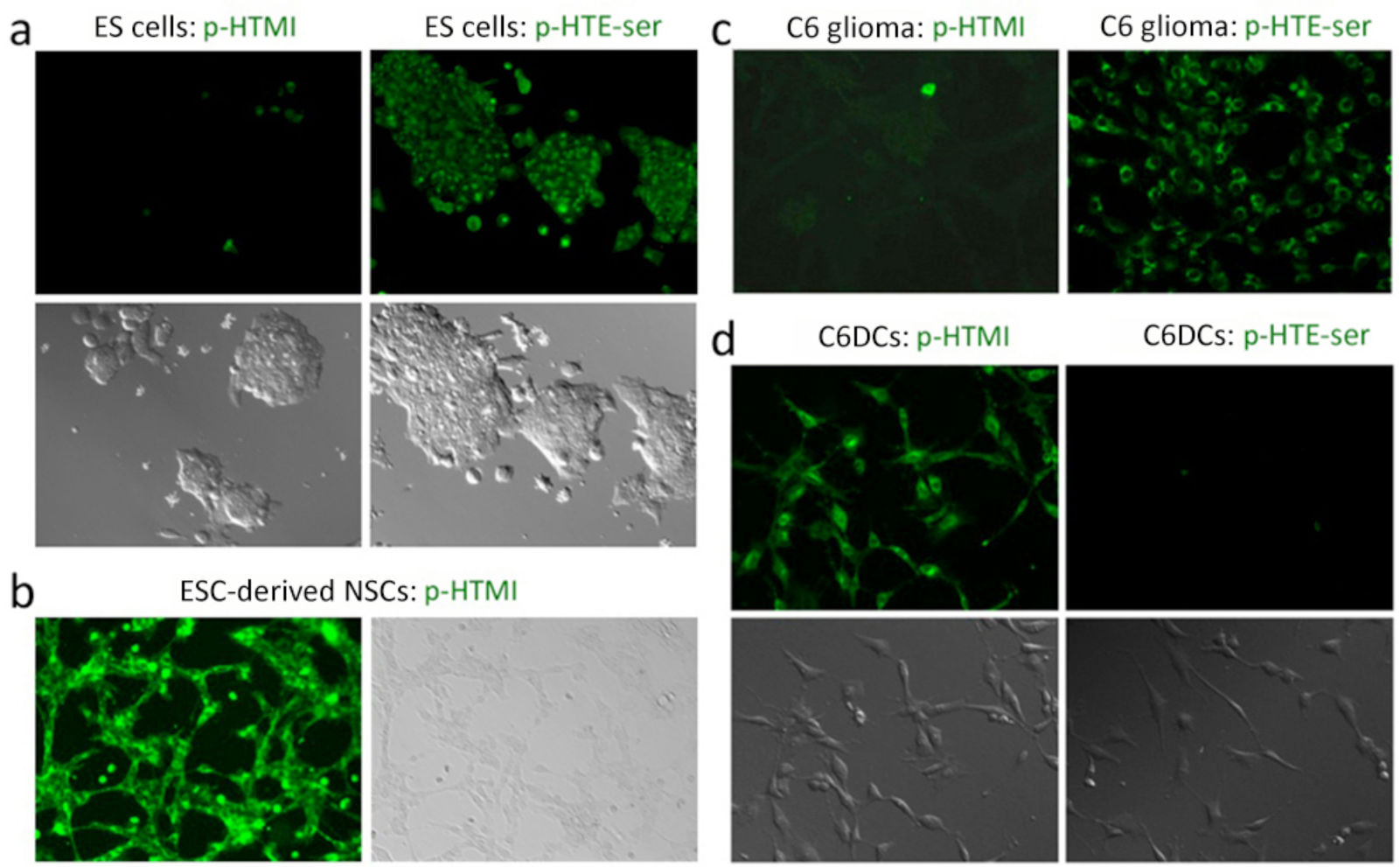
p-HTMI detects C6 glioma-derived and embryonic stem cell derived NSCs. (**a**) Very few (≪1%) ESCs display staining by p-HTMI, whereas p-HTE-Ser stains ≫90% of ESCs. (**b**) p-HTMI stains virtually 100% of ES cell-derived NSCs. (**c**) Only small subsets of conventionally cultured C6 glioma cells with distinct morphology are stained by p-HTMI, whereas p-HTE-Ser stains ≫90% of these cells. (**d**) Basically 100% of FGF2-exposed C6 glioma derived cells (C6DCs) are stained by p-HTMI, whereas p-HTE-ser does not stain these cells.

Glioblastoma (GBM) is a common and aggressive type of brain tumor in adult humans [4]. Rat C6 glioma can be considered as an experimental model system for studying GBM. C6 glioma cells do not normally respond efficiently to differentiation-inducing factors, which was confirmed by our experiments (**Figure 3a**). We therefore applied the protocol for embryonic rat NSCs to C6 glioma cells to see if the differentiation potential could be affected. Grown in the presence of FGF2 on fibronectin coated plates and in N2 medium with supplements, C6 glioma cells grown as NSCs (referred to as C6DCs) formed uniform cultures of nestin-positive cells (**Figure 3b**). In the C6DCs treated with CNTF, clear signs of increased astrocytic differentiation were detected (**Figure 3b**), and valproic acid (VPA) induced differentiation into neuronal-like, TuJ1-positive cells (**Figure 3b**) as previously shown for embryonic rat NSCs [14]. In addition, the gene expression of *gfap* and *Tubb3* was elevated upon CNTF and VPA treatment, respectively (**Figure 3c**). Thus, rat C6 glioma cells grown in similar conditions as embryonic rat NSCs become responsive to external factors and obtain at least a subset of characteristics of neural stem cells.

**Figure 3.**
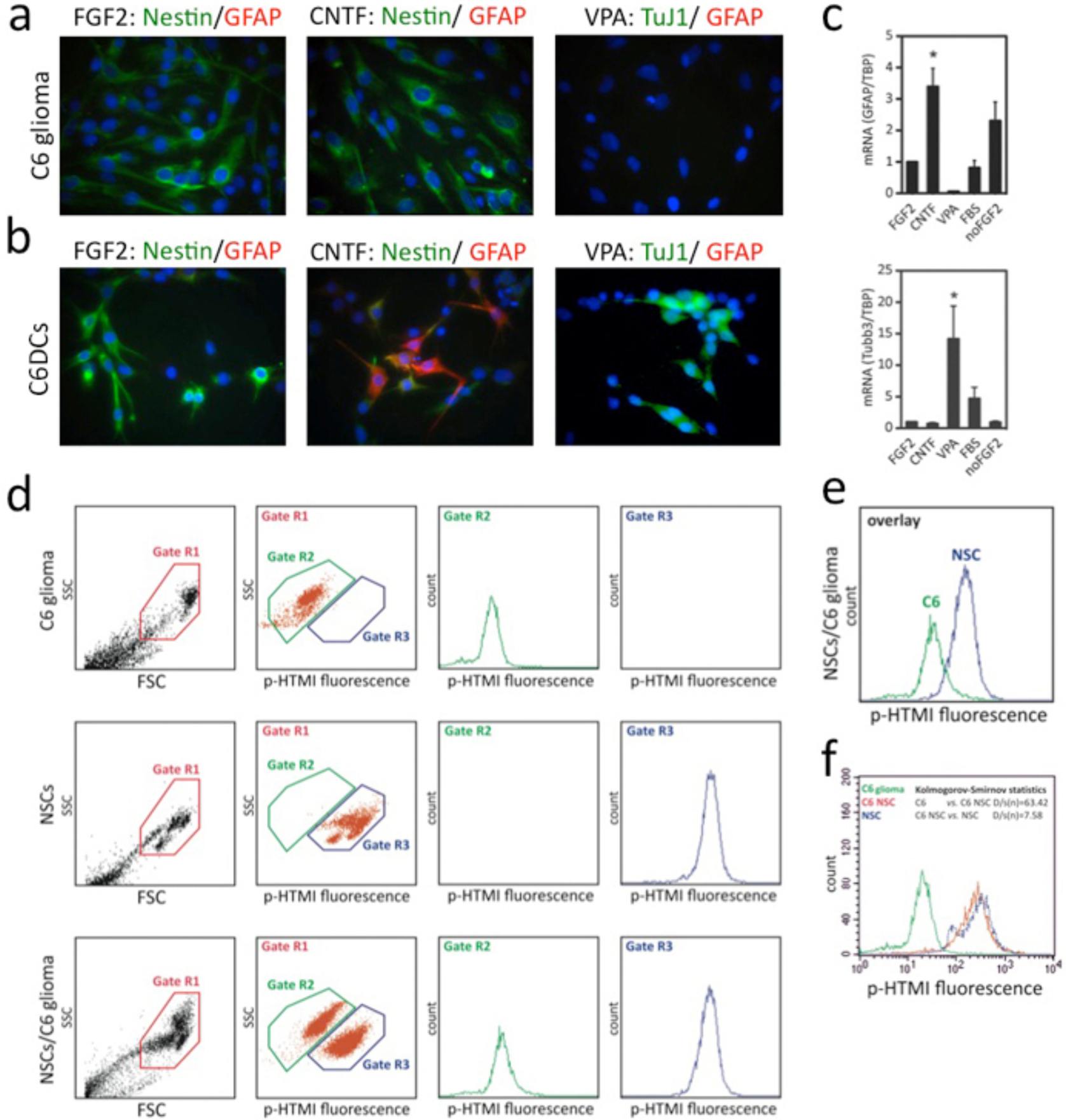
p-HTMI specifically detects glioma-derived stem-like cells by FACS. (**a**) Conventionally cultured C6 glioma cells are Nestin-positive but stain negative for differentiation markers, also after treatment with CNTF or VPA. (**b,c**) C6 glioma cells cultured in 2D layers in the presence of FGF2 respond to CNTF and VPA and increased numbers of GFAP-positive cells and TuJ1-positive cells (**b**), along with increased mRNA levels (**c**), are detected after administration of CNTF and VPA, respectively. (**d,e,f**) Cell sorting by FACS demonstrates that p-HTMI specifically and selectively detects glioma-derived stem-like cells and neural stem cells but not glioma cells.

Various studies have suggested that 1-4% of the conventionally cultured C6 glioma cells share characteristics of cancer stem cells [15]. We administered p-HTMI to regularly cultured C6 glioma cells and noted that this resulted in a staining of around 1-2% of the cells (**Figure 2c**). In contrast, administration of p-HTE-Ser resulted in staining of >95% of the C6 glioma cells (**Figure 2c**). We next stained C6DCs grown in the presence of FGF2 using p-HTMI or p-HTE-Ser. Surprisingly, the C6DCs stained with p-HTMI now displayed a strong fluorescent signal, while p-HTE-Ser did not stain any cells **(Figure 2d**). FACS analysis confirmed these results and verified that a population of approximately 1% of regular C6 glioma cells is detected by p-HTMI (**Figure 3d,e**). This experiment further validated the ability of p-HTMI to distinguish between cell types, as the vast majority of C6DCs and embryonic NSCs could be clearly distinguished from conventionally cultured C6 glioma cells (**Figure 3f**).

As mentioned above, an advantage with previously developed polythiophene derivatives is that these have been shown to be applicable *in vivo* in patient material, for example when detecting amyloid aggregates in tissue from patients with Alzheimer’s Disease [2b]. An *in vivo* application for p-HTMI could be of clinical importance for future development of use in surgical resection of GBM. To investigate whether p-HTMI could be applied also in an *in vivo* context, we transplanted either cells of a classic human GBM cell line (U-87MG; [16]) or cells derived more recently from GBM patients [17] into the right striatum of mice as previously described [16]. The animals were monitored on a daily basis for neurological symptoms associated with tumor growth. The animals were sacrificed 5-7 weeks after transplantation of U-87MG and 16-24 weeks after transplantation of human GBM cells. Delivery of a terminal dose of avertin was followed by an injection of p-HTMI into the original site of glioma cell transplantation. The brains were then removed, cut by microtome, and the sections were investigated histologically in the microscope. The initial experiments using U-87MG cells did reveal a small (1-5%) population of human cells stained by injected p-HTMI (data not shown). Analysis of the mice with tumors derived from cells from GBM patients [17] revealed that p-HTMI indeed detected a subpopulation of human cells in sections from the tumor and surrounding brain tissue (**Figure 4c-d, Supplementary Movie 2**) whereas non-injected tumor tissue remained unstained (**Figure 4a-b**), in spite of occasional presence of p-HTMI in the perfusion solution (see **Methods**). The results suggest that p-HTMI can be applied in an *in vivo* context by direct injection into the tumor. While tumor-free brain tissue remained unlabeled, we did notice labeling of extracellular debris, dead cells and also murine cells, especially around the injection site and within or in close proximity to the tumor (**Figure 4c-d**).

**Figure 4.**
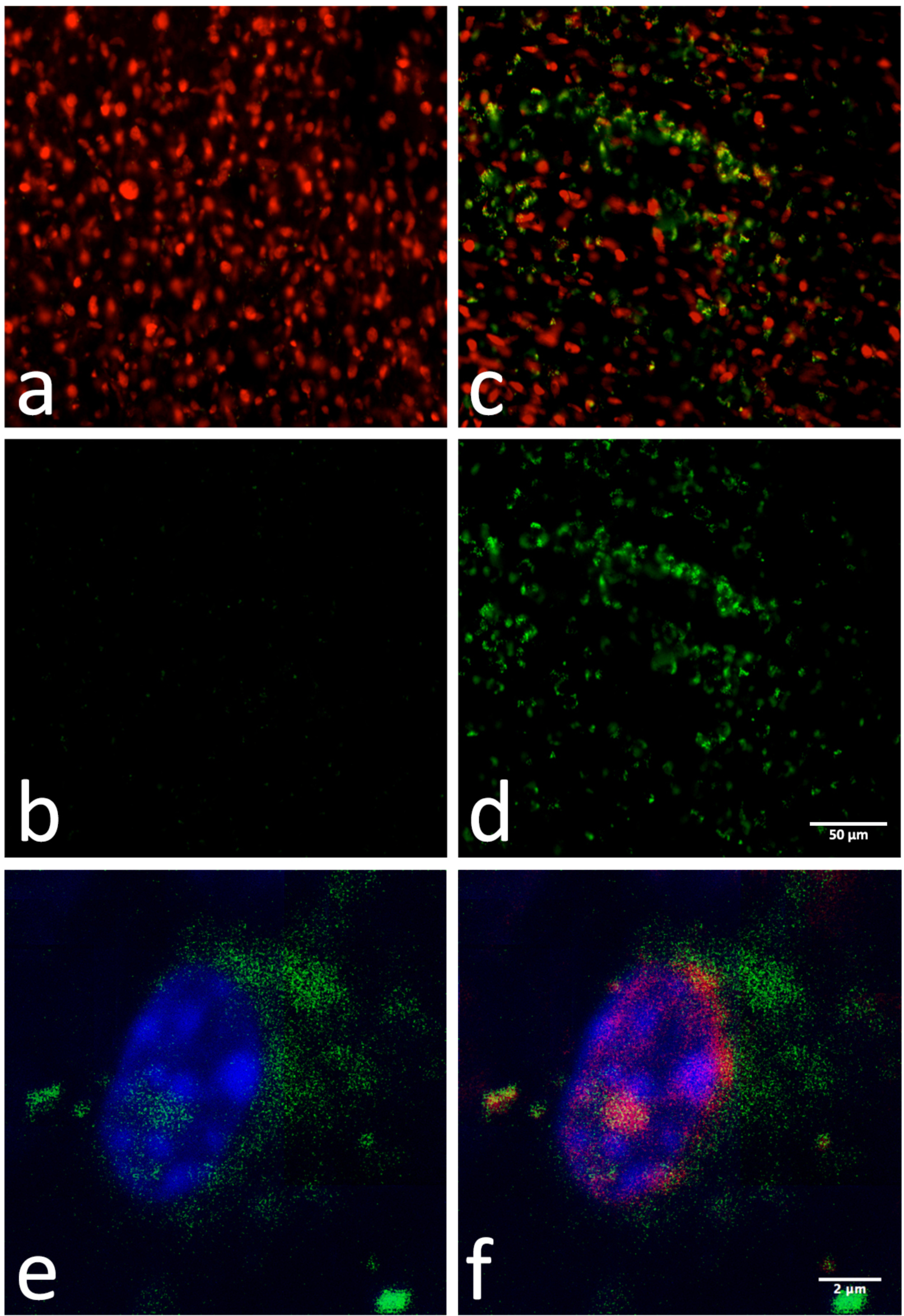
p-HTMI detects a subpopulation of human cells in brain sections with tumors derived from patients with GBM. (**a-d**) Micrographs depicting brain sections from NOD/SCID mice injected with GBM cells (U3031MG) and injected with either vehicle (a,b) or p-HTMI (c,d) prior to perfusion. P-HTMI-positive cells depicted in green and human tumor cells stained with human nuclei antibody depicted in red (20x). Scale bar approximately 50 µm. (**e-f**) Micrographs from a confocal microscope demonstrating cytoplasmic staining of p-HTMI in a human GBM cell from a mouse brain section. Green is p-HTMI signal, red is the human nuclei antibody, and blue is DAPI. Please note that the blue and red channels were partially cut in figure e and f as a neighboring human cell nucleus, negative for p-HTMI, obscured the signal. See also Supplementary Movie 2.

The obtained results from the C6 glioma cell line *in vitro* and the orthotopic tumors *in vivo* prompted an investigation of whereas the human GBM cells detected by p-HTMI shared characteristics of so called GSCs. We therefore used three previously characterized cell lines (U3034MG, U3088MG, U3031MG) derived from human GBM expanded *in vitro* [17] from different individual patients with glioblastoma. Such patient-derived cells has been shown to contain a high proportion of tumor initiating cells (TICs), to stain positive for SOX2 and nestin, and have the ability to self-renew as well as differentiate into various neural cell fates [17]. FACS experiments demonstrated that p-HTMI reproducibly detected between 70-90% of the cells in these three cultures within 10 minutes from application of the molecule (**Figure 5b,c,e**). To assess whether p-HTMI exerted any effect on the cell survival of these cells, we investigated the cell death in the cultures of the treated cells, but found no significant difference between control cultures and p-HTMI-treated cells when analyzing PI-staining and the number of Annexin V-positive cells by FACS (**Figure 5a**).

**Figure 5.**
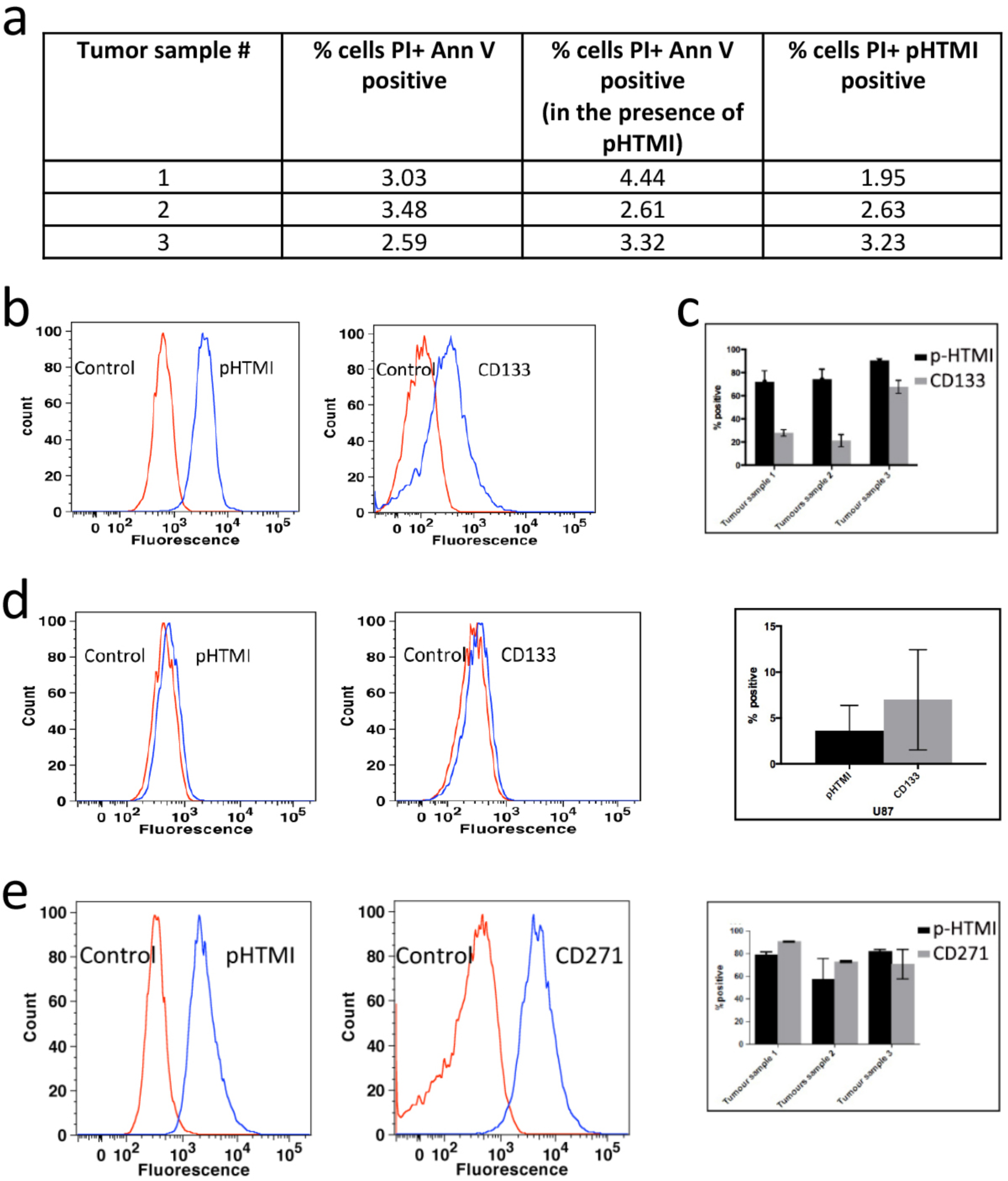
p-HTMI detects the same population of GBM cells as CD271. (**a**) Minimal cell necrosis and apoptosis in all three tumor samples in the presence of p-HTMI verified using Propidium Iodide (PI) and Annexin V. (**b,c**) Comparison of p-HTMI and CD133-labeled GBM cells as assessed by fluorescence-activated cell sorting (FACS). (**d**) A small number of cells were detected by p-HTMI and CD133 in U-87MG glioma cells. (**e**) In a double-labeled cell population (CD271+p-HTMI), p-HTMI and CD271 label largely the same population of cells in U3034MG, U3088MG, and U3031MG nestin+/SOX2+ patient-derived GBM cells.

We next aimed at elucidating the population of p-HTMI-positive cells detected in the GC cultures by comparison with antibodies previously discussed to detect stem-like cells from various sources of cancer. CD133 or prominin is an antigen expressed by various neural progenitor populations and it has been shown that depletion of CD133-positive cells from glioma cell populations decreases, but does not abolish, the tumor-initiating ability of transplanted glioma cells [4]. In the GC cultures examined, CD133 stained 20-70% of the cells in comparison to p-HTMI that stained 70-90% (**Figure 5b,c**). To investigate whether the higher number of cells detected by p-HTMI was merely the result of a less specific labeling, we compared labeling of CD133 antibody and p-HTMI in the tumor cell line U-87MG, previously shown to contain a very small percentage of GSCs in cell culture [16]. The number of cells detected by CD133 antibody as well as p-HTMI was much lower in the U-87MG cultures, and in these cells, the number of cells detected by CD133 antibody was higher than that of p-HTMI (**Figure 5d**). CD44 antibodies are under thorough investigation in various cancers for their ability to detect progenitor cells *in vivo*, but may not be suitable for use *in vitro*. Here, CD44 antibody detected basically all cells (>98%) in the three GC cultures as well as the U-87MG cell line and thus showed significantly less specificity than p-HTMI (**Supplementary Figure S4, Figure 5b,c,e, and data not shown**).

CD271, or nerve growth factor receptor p75, has been reported to be expressed by various types of progenitor cells in many organs and in different tumor types. In accordance, it has been shown that CD271 antibody selectively detects neural progenitors *in vitro* and *in vivo* [18]. Intriguingly, it has been shown that CD271 antibody detects two subpopulations of cells in GBM, stem cell-like cells and the rapidly migrating cells [19]. We therefore pursued a double staining experiment with p-HTMI and CD271 antibody in GC cultures, and found by FACS that the cell populations in GC cultures stained by p-HTMI and CD271 are vastly overlapping (**Figure 5e**). We conclude that p-HTMI-positive cells in GCs are similar to those labeled by CD271 and thus p-HTMI-stained cells may be subpopulations of stem cell-like cells and/or rapidly migrating GBM cells.

Here we have demonstrated that p-HTMI is a novel molecular probe that specifically stains live progenitors derived from embryonic rodent brains and rodent and human glioma within 10 minutes by just mere application of the molecule to the cell culture or injection into a tumor, and is thus representative of a new generation of smart molecules to be used for non-invasive, non-genetic live cell detection. It should be noted that more studies are required to determine the true character of the human cells detected by p-HTMI *in vivo*. Initial attempts to label the p-HTMI-positive cells with CD133 antibody resulted in a large proportion of double-labeled cells (B.M., O.H., unpublished observations), but the validity of CD133 antibodies in immunohistochemical applications have been questioned, and thus isolation of p-HTMI-positive cells followed by single cell analysis is required to thoroughly investigate whether the p-HTMI-positive population is homogenous or not. In addition, it will be of immediate interest to pursue *in vivo* studies in animals to investigate the possibility to use p-HTMI as a complementary tool to detect GSCs in fluorescence-guided surgical resection of GBM. CD271 antibody stains for cells derived from and in various organs and is significantly less cell-specific than p-HTMI, and further provides only an indirect detection of the cells. The result thus strengthens the verification of the specificity and versatility of p-HTMI. Future studies will aim at in detail characterization of the GBM cells detected by p-HTMI as well as the intracellular target of the molecule.

## Acknowledgements

We thank Drs Anna Herland, Vanessa Lundin, and Johan Holmberg for valuable discussions and cells, and Olle Inganäs for support. This work was supported by the Swedish Cancer Society (CF), the Swedish Research Council (VR-M), the K&A Wallenberg Foundation (CLICK), the Swedish Foundation for Strategic Research (SSF; OBOE project), European Research Council (ERC; Project MUMID), and the Swedish Childhood Cancer Foundation (BCF).

## Conflict of interest

S.I., R.S., A.K.O.Å., P.K., O.H., and K.P.R.N. are co-inventors on a patent application regarding the structure and applications of p-HTMI described in the manuscript.

## Methods (Supplementary)

### Synthesis

#### General Methods

**Scheme 1.**
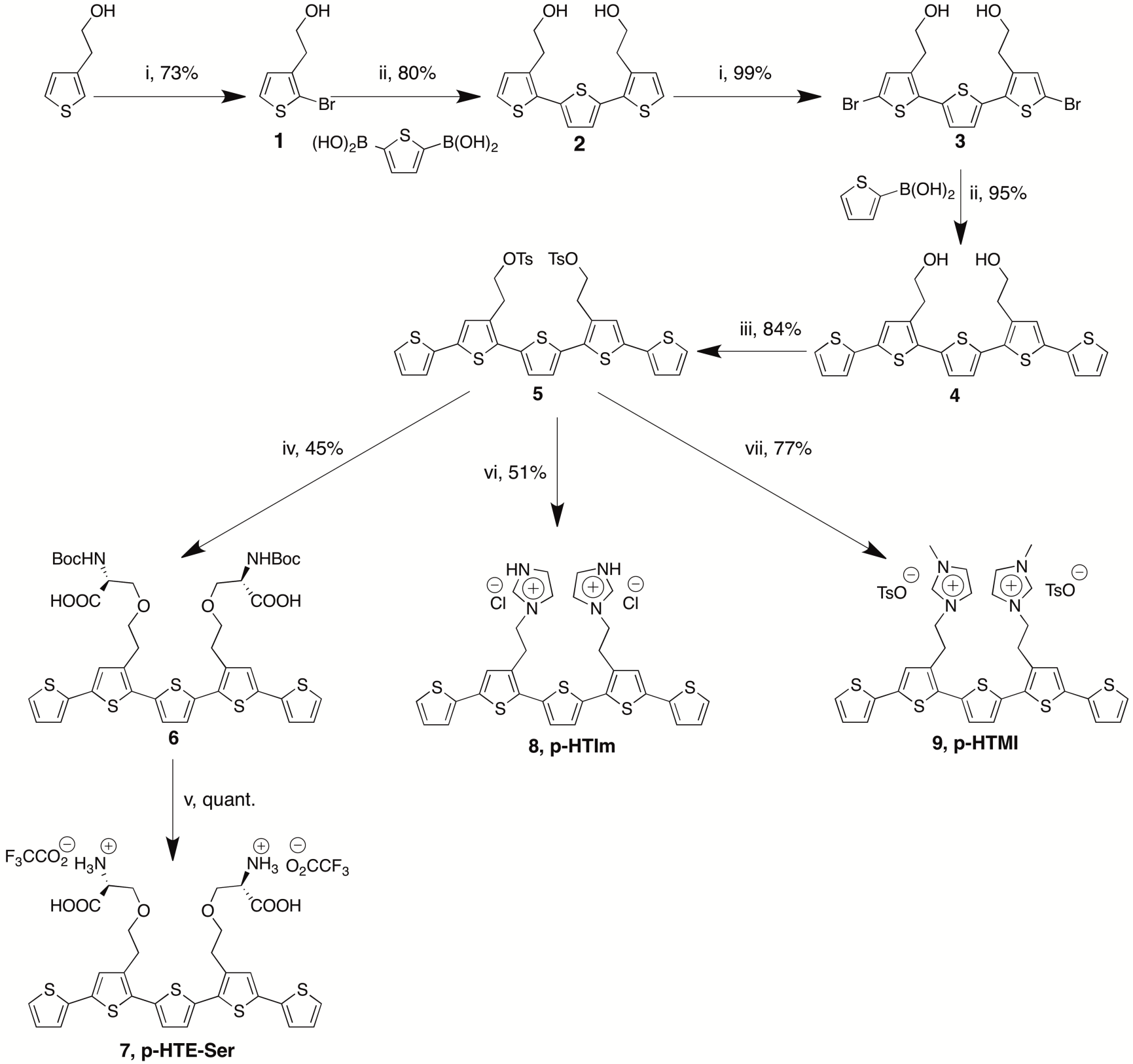
Reagents and conditions: (i) *N*-bromosuccinimide, DMF, −15 °C; (ii) PEPPSI-IPr, K_2_CO_3_, toluene/methanol (1:1), 75 °C; (iii) *p*-toluenesulphonyl chloride, CHCl_3_, pyridine; (iv) methylimidazole, acetonitrile, 75 °C; (v) imidazole, acetonitrile; (vi) Boc-L-Ser-OH, K_2_CO_3_, DMF, 50 °C; (vii) TFA, CH_2_Cl_2_.

Organic extracts were dried over anhydrous magnesium sulfate, filtered and concentrated *in vacuo* at 40 °C. 3-thiopheneethanol, 2-thiopheneboronic acid, thiophene-2,5-bisboronic acid, and PEPPSI-IPr([1,3-Bis(2,6-Diisopropylphenyl)imidazol-2-ylidene](3-chloropyridyl)palladium(II) dichloride) are commercially available from Sigma-Aldrich Co. NMR-spectra were recorded on a Varian instrument (^1^H 300 MHz, ^13^C 75.4 MHz). Chemical shift were assigned with the solvent residual peak as a reference according to Gottlieb *et al*. [20]. TLC was carried out on Merck precoated 60 F254 plates using UV-light (λ = 254 nm and 366 nm) and charring with ethanol/sulfuric acid/*p-*anisaldehyde/acetic acid 90:3:2:1 for visualization. Flash column chromatography (FC) was performed using silica gel 60 (0.040-0.063 mm, Merck). Gradient HPLC-MS was performed on a Gilson system (Column: Waters X-Bridge C-18 or C-8 5 µ, 250 × 15 mm and Waters X-Bridge C-18 2.5 µ, 150 × 4.6 mm for semipreparative and analytical runs respectively; Pump: Gilson gradient pump 322; UV/VIS-detector: Gilson 155; MS detector: Thermo Finnigan Surveyor MSQ; Gilson Fraction Collector FC204) using acetonitrile with 0.05% formic acid or triethylamine and deionized water with 0.05% formic acid or triethylamine as mobile phase. For preparative reversed phase purifications a VersaFlash™ system equipped with VersaPak™ C18 cartridges. MALDI-TOF MS was recorded in linear positive mode with α-cyano-4-hydroxycinnamic acid matrix (CHCA) or 2,5-dihydroxy benzoic acid (DHB) as matrix.

#### Synthesis of 1

3-thiopheneethanol (20.19 g, 157.5 mmol) was dissolved in DMF (100 mL) and the solution was cooled to −15 °C. NBS (22.4 g, 125.9 mmol) was added portion wise during one minute. The solution was allowed to attain room temperature during 2 h, again cooled to −15 °C, more NBS (5.65 g, 31.7 mmol) was added portion wise during one minute, and the solution was allowed to attain room temperature. The reaction mixture was diluted with EtOAc, washed with brine, dried, filtered and concentrated. Purification by FC (DCM) and reversed phase VersaFlash™ gave **1** (73%) as colorless oil. R_*f*_: 0.31 (toluene/ethyl acetate 4:1). ^1^H NMR (CDCl_3_) δ 2.84 (t, 2H, *J* = 6.6 Hz), 3.81 (t, 2H, *J* = 6.6 Hz), 6.82 (d, 1H, *J* = 5.2 Hz), 7.41 (d, 1H, *J* = 5.2 Hz); ^13^C NMR (CDCl_3_) δ 35.5, 62.4, 75.6, 128.5, 131.0, 143.3.

#### Synthesis of 2

A mixture of **1** (11.8 g, 57.0 mmol), 2.5-thiophenediboronic acid (4.80 g, 27.9 mmol), and K_2_CO_3_ (18.0 g, 130.2 mmol) in MeOH/toluene 1:1 (150 mL) was heated at 75 °C for 1 min. PEPPSI™-IPr catalyst (0.190 g, 0.280 mmol) was added and the mixture was heated at 75 °C for an additional 30 min. HOAc (conc.) and EtOAc were added and the mixture was washed with brine and water. The organic phase was dried, filtered and concentrated. Purification by reversed phase VersaFlash™ gave **2** (80%) as slightly yellow oil. R_*f*_: 0.17 (toluene/ethyl acetate 2:1) Calcd mass for C_16_H_16_O_2_S_3_: [M+H]^+^ 337.0; found 336.9. [M+Na]^+^ 359.0; found 359.0. ^1^H NMR (CDCl_3_) δ 3.07 (t, 4H, *J* = 6.9 Hz), 3.89 (t, 4H, *J* = 6.9 Hz), 6.99 (d, 2H, *J* = 5.2 Hz), 7.10 (s, 2H), 7.22 (d, 2H, *J* = 5.2 Hz); ^13^C NMR (CDCl_3_) δ 32.7, 63.0, 124.6, 126.8, 130.2, 132.1, 135.4, 135.9.

#### Synthesis of 3

**2** (328 mg, 0.975 mmol) was dissolved in DMF (3 mL) and the solution was cooled to −15 °C. NBS (351 mg, 1.97 mmol) was added portion wise during one minute. The solution was allowed to attain room temperature during 2h. DCM and brine were added and the organic phase was separated, washed twice with 1M HCl, dried, filtered, and concentrated. The slightly crude yellow product **3** (99%) was used without further purification in the next step. ^1^H NMR (300 MHz, CDCl_3_/CD_3_OD 1:1) δ: 2.92 (t, *J* = 7.0 Hz, 4H), 3.74 (t, *J* = 7.0 Hz, 4H), 6.95 (s, 2H), 7.00 (s, 2H); ^13^C NMR (75.5 MHz, CDCl_3_/CD_3_OD 1:1) δ: 32.7, 62.2, 111.4, 127.4, 133.2, 133.4, 135.4, 137.1.

#### Synthesis of 4

To a solution of **3** (463 mg, 0.937 mmol) in MeOH/toluene 1:1 (5 mL) were added thiophene-2-boronic acid (475 mg, 3.71 mmol), and K_2_CO_3_ (647 mg, 4.68 mmol). The mixture was heated at 75 °C for 1 min and PEPPSI™-IPr catalyst (14 mg, 0.021 mmol) was added and the mixture was heated at 75 °C for an additional 20 min. HOAc (conc.) and EtOAc were added and the mixture was washed with brine and water. The organic phase was dried, filtered and concentrated. Short FC (ethyl acetate) gave product **4** (95%) as orange solid. ^1^H NMR (300 MHz, DMSO-d6) δ 2.91 (t, *J* = 6.8 Hz, 4H), 3.68-3.77 (m, 4H), 4.83 (t, *J* = 5.2 Hz, 2H), 7.06-7.12 (m, 2H), 7.27 (s, 2H), 7.31 (d, *J* = 3.4 Hz, 2H), 7.51 (d, *J* = 5.0 Hz, 2H); ^13^C NMR (75.5 MHz, DMSO-d6) δ 32.6, 60.7, 124.2, 125.7, 126.8, 127.4, 128.4, 129.0, 134.5, 134.8, 135.9, 137.7.

#### Synthesis of 5

Product **4** (0.049 g, 0.0979 mmol) and *p*-toluenesulphonyl chloride (0.056 g, 0.294 mmol) were added to a mixture of CHCl_3_ (0.8 mL) and pyridine (0.2 mL). After 4h the solution was diluted with toluene and washed with HCl (1 M, aq.) and H_2_O. FC (toluene) gave product **5** (84%) as yellow solid. ^1^H NMR (300 MHz, CDCl_3_) δ 2.32 (s, 6H), 3.11 (t, *J* = 6.6 Hz, 4H), 4.28 (t, *J* = 6.6 Hz, 4H), 6.86 (s, 2H), 7.01 (s, 2H), 7.03 (dd, *J* = 3.6, 5.1 Hz, 2H), 7.14 (dd, *J* = 1.1, 3.6 Hz, 2H), 7.20-7.26 (m, 6H), 7.69 (d, *J* = 8.3 Hz, 4H); ^13^C NMR (75.5 MHz, CDCl_3_) δ 21.6, 29.0, 69.4, 124.1, 125.0, 126.2, 127.0, 127.9, 128.1, 129.9, 131.0, 132.8, 134.0, 135.3, 136.5, 136.6, 144.9.

#### Synthesis of 6

(0.120 g, 0.148 mmol), Boc-L-Ser-OH (0.122 g, 0.593 mmol) and potassium carbonate (0.122 g, 0.890 mmol) were dissolved in 3 mL DMF and heated to 50 °C. After 1 day the reaction mixture was diluted with ethyl acetate and washed twice with HCl (1M, aq.) and three times with brine. The crude product was purified on FC (toluene/ethyl acetate 4:1) followed by HPLC ACN/H_2_O 8:1->1:0 over 30 minutes to give product **6** in 45% yield as an orange solid. TLC (toluene/ethyl acetate 1:1) R_*f*_: 0.30. ^1^H NMR (300 MHz, CDCl_3_) δ 1.43 (s, 18H), 3.17 (t, *J* = 7.1 Hz, 4H), 3.85 (dd, *J* = 3.6, 11.2 Hz, 2H), 3.92 (dd*J* = 3.7, 11.2 Hz, 2H), 4.37-4.50 (m, 6H), 5.51 (d, *J* = 5.8 Hz, 2H), 7.02 (dd, *J* = 3.6, 5.1 Hz, 2H), 7.05 (s, 2H), 7.13 (s, 2H), 7.18 (dd, *J* = 1.1, 3.6 Hz, 2H), 7.23 (dd, *J* = 1.1, 5.1 Hz, 2H); ^13^C NMR (75.5 MHz, CDCl_3_) δ 28.4, 28.8, 56.0, 63.6, 65.1, 80.5, 124.2, 125.0, 126.4, 126.9, 128.3, 131.1, 134.7, 135.5, 136.4, 136.7, 155.9, 171.0.

#### Synthesis of p-HTE-Ser (7)

**6** (0.048 g, 0.055 mmol) was dissolved in dichloromethane/TFA (4:1, 5 mL). After 2 h the solution was co-concentrated with toluene to give product **7** as red solid quantitively. ^1^H NMR (300 MHz, DMSO-d6) δ 3.12 (t, 4H, *J* = 6.6 Hz), 3.69 (dd, 2H, *J* = 3.9, 11.6 Hz), 3.74 (dd, 2H, J = 4.3, 11.6 Hz), 3.93 (t, 3.9 Hz, 2H), 4.44 (t, *J* = 6.6 Hz, 4H), 7.11 (dd, *J* = 3.6, 5.1 Hz, 2H), 7.30 (s, 2H), 7.34 (dd, *J* = 1.1, 3.6 Hz, 2H), 7.35 (s, 2H), 7.55 (dd, *J* = 1.1, 5.1 Hz, 2H); ^13^C NMR (75.5 MHz, DMSO-d6) δ 28.0, 54.8, 60.5, 64.6, 124.5, 126.0, 127.0, 127.3, 128.5, 129.4, 134.5, 135.2, 135.6, 135.7, 169.5

#### Synthesis of p-HTIm (8)

**5** (116 mg, 0.143 mmol) and imidazole (500 mg, 7.34 mmol) were dissolved in MeCN (1.5 mL). The solution was heated at 65 °C for 2h and concentrated. FC (EtOAc + TEA and EtOAc/MeOH 4:1 + 1% TEA) gave crude compound that was further purified by gradient HPLC to provide p-HTIm (**8**) (51%) as orange solid. NMR was run on neutral compound, after which the sample was dissolved in 1M HCl, concentrated and dried to give a red-brown chloride salt. ^1^H NMR (300 MHz, CD_3_OD) δ 3.18 (t, *J* = 6.8 Hz, 4H), 4.27 (t, *J* = 6.8 Hz, 4H), 6.90 (app. t, *J* = 1.2 Hz, 2H), 6.92 (s, 2H), 6.93 (s, 2H), 6.97 (app. t, *J* = 1.3 Hz, 2H), 7.01 (dd, *J* = 3.6, 5.1 Hz, 2H), 7.18 (dd, *J* = 1.1, 3.6 Hz, 2H), 7.33 (dd, *J* = 1.1, 5.1 Hz, 2H), 7.46 (app. t, *J* = 1.1 Hz, 2H); ^13^C NMR (75.5 MHz, CD_3_OD) δ 31.9, 48.2, 120.7, 125.2, 126.2, 127.2 128.1, 129.1, 129.1, 131.8, 136.3, 137.0, 137.6, 137.8, 138.3.

#### Synthesis of p-HTMI (9)

**5** (1.00 g, 1.24 mmol) was dissolved methylimidazole (4.75 mL) and DMF (1 mL). The mixture was heated at 75 °C for 4 hours and concentrated and co-concentrated with xylene to remove excess methylimidazole. Purification by gradient HPLC gave p-HTMI (**9**) (77%) as orange tosylate/formic acid salt. LC-MS calcd for C_32_H_30_N_4_S_5_: [M]^2+^: 315.05; Found 315.10. ^1^H NMR (300 MHz, D_2_O, 35 °C) δ 2.22 (s, 4.4H, tosylate), 3.20 (t, *J* = 7.4 Hz, 4H), 3.91 (s, 6H), 4.33 (t, *J* = 7.4 Hz 4H), 7.05 (d, *J* = 4.2, 2H), 7.07 (s, 4H), 7,12 (d, *J* = 8.0 Hz, 2.9H, tosylate), 7.2 (d, *J* = 3.5 Hz, 2H), 7.28 (s, 2H), 7.35 (d, *J* = 5.1 Hz), 7,45 (s, 2H), 7.70 (d, *J* = 8.0 Hz, 3H, tosylate), 8.62 (s, 0.8H, formic acid), 8.66 (s, 2H); ^13^C NMR (75.5 MHz, D_2_O, 35 °C) δ 20.9, 29.6, 36.1, 49.1, 122.2, 124.0, 124.6, 125.7, 125.9, 126.3, 127.5, 128.8, 129.3, 130.8, 134.5, 134.9, 136.1, 136.3, 136.6, 141.4, 141.4.

**Scheme 2.**
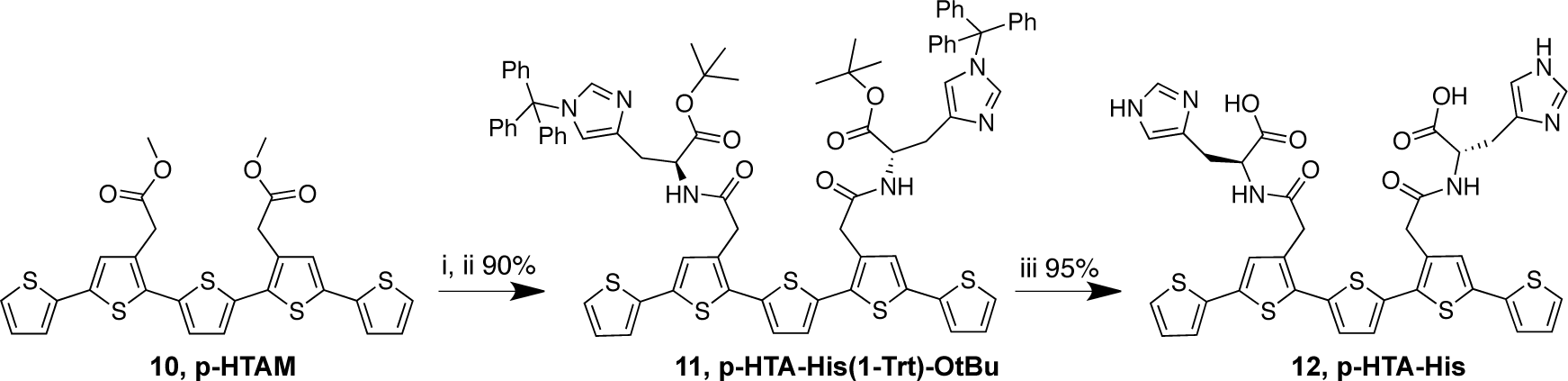
Reagents and conditions: (i) 1M NaOH, dioxane, H_2_O; (ii) H-His(1-Trt)-OtBu, DIPEA, *O*-(7-azabenzotriazol-1-yl)-*N,N,N’,N’-*tetramethyluronium hexafluorophosphate (HATU), DMF; (iii) Et_3_SiH, TFA, DCM.

#### Synthesis of p-HTA-His(1-Trt)-OtBu (11)

To a solution of p-HTAM (**10**) (93 mg, 0.167 mmol) in dioxane (2 mL) and H_2_O (0.5 mL) was added 1M NaOH (0.340 mL, 0.340 mmol). The solution was heated at 65 °C for 1 h, neutralized with 1M HCl, and the solvents were co-evaporated with toluene. The residual was dissolved in DMF (3 mL) and H-His(1-Trt)-OtBu (302 mg, 0.666 mmol) and DIPEA (0.100 mL, 1.04 mmol) were added. The temperature of the solution was lowered to 0 °C and *O*-(7-azabenzotriazol-1-yl)-*N,N,N’,N’-* tetramethyluronium hexafluorophosphate (HATU) (190 mg, 0.500 mmol) was added. The solution was allowed to stir for 30 min at 0 °C and then for 4 h at room temperature. EtOAc, toluene and brine were added and the organic phase was separated and washed with saturated NH_4_Cl (aq), saturated NaHCO_3_ (aq) and H_2_O, dried, filtered and concentrated. FC (toluene/EtOAc 1:1 + 1 % TEA gave p-HTA-His(1-Trt)-OtBu (**11**) (90 %) as yellowish oil. ^1^H NMR (300 MHz, CDCl_3_) δ 1.32 (s, 18H), 2.86- 3.01 (m, 4H), 3.64 (d, *J* = 16.5 Hz, 2H), 3.70 (d, *J* = 16.5 Hz, 2H), 4.60-4.69 (m, 2H), 6.49 (d, *J* = 1.2 Hz, 2H), 6.92 (dd, *J* = 3.6, 5,1 Hz, 2H), 6.98-7.04 (m, 12H), 7.05 (dd, *J* = 1.1, 3.6 Hz, 2H), 7.12 (dd, *J* = 1.1, 5.1 Hz, 2H), 7.14-7.17 (m, 6H), 7.20-7.35 (m, 18H), 7.78-7.84 (b, 2H); ^13^C NMR (75.5 MHz, CDCl_3_) δ 28.2, 29.5, 37.4, 53.3, 75.3, 81.3, 119.3, 123.9, 124.7, 127.6, 127.9, 128.1, 129.9, 132.1, 132.2, 135.5, 136.1, 136.6, 137.0, 138.6, 142.4, 169.9, 170.3.

#### Synthesis of p-HTA-His (12)

pHTA-His(1-Trt)-OtBu (**11**) (35 mg, 0.025 mmol) was dissolved in DCM (1 mL) and Et_3_SiH (0.035 mL, 0.414 mmol) was added. TFA (1 mL) was added and the solution was stirred for 3 h. The completeness of the reaction was validated by HPLC-MS. Solvents were co-evaporated with toluene. Purification by HPLC-MS gave p-HTA-His (**12**) (95 %) as orange solid. ^1^H NMR (300 MHz, (CD_3_)_2_SO) δ 2.88-3.14 (m, 4H), 3.62 (s, 4H), 4.45-4.60 (m, 2H), 7.07 (s, 2H), 7.11 (dd, *J* = 3.6, 5.1 Hz, 2H), 7.23 (s, 2H), 7.29 (s, 2H), 7.30 (dd, *J* = 1.1, 3.6 Hz, 2H), 7.55 (dd, *J* = 1.1, 5.1 Hz, 2H), 8.13 (s, 2H), 8.52-8.59 (b, 2H).

### Cell culture

Rat embryonic neural stem cells (NSCs) and mouse embryonic stem cells were derived and cultured as previously described (Johe, Hazel et al. 1996, Andersson, Duckworth et al. 2011). Briefly, NSCs were harvested from the cerebral cortices of E15.5 embryos of timed pregnant Sprague-Dawley rats. Cells were mechanically dissociated and plated on previously coated tissue culture plates. NSCs and C6 (ATCC CCL-107) -derived stem cell-like cells were expanded in N2 medium supplemented with 10 ng/ml of FGF2 (R&D Systems) until 80% confluency. Animals were treated in accordance with institutional and national guidelines (ethical permits no. N310/05; N79/08; N284/11).

### Staining of cells

In general, the LCOs were dissolved in deionized water to a final concentration of 1 mg/ml, and administered at a dilution of 1:500 (1:10-1:10 000 were tested), directly to each well and the detection was done after 10 min in a fluorescent microscope or FACS. p-HTMI generated fluorescence at a wavelength common to green fluorescent proteins.

### Two-photon microscopy of NSCs stained with p-HTMI

The emission wavelength of p-HTMI was characterized in a two-photon microscope. NSCs were grown in 35 mm plates (40 000 cells/plate) and treated with 10 ng/ml FGF2 for 48 hours. Prior to staining the cells, the medium was changed to DMEM:F12 medium without phenol red (Invitrogen) in order to eliminate background signals. CellTracker (Invitrogen) was administered together with p-HTMI (1:500) in order to be able to find living cells, resulting in a double staining in red (CellTracker) and green (p-HTMI).

### Immunocytochemistry

The plates were first rinsed once in PBS and then fixed in 10% formaldehyde for 20 min. The formaldehyde was aspirated and the plates were washed 3 times, 5 min each, in PBS/0.1% Triton-X 100. The plates were then incubated with respective primary antibody in PBS/0.1% Triton-X 100/ 1% BSA overnight at 4 °C. The samples were then washed 6 times, 5 min each, in PBS/0.1% Triton-X 100. Secondary antibodies (1:500) in PBS/0.1% Triton-X 100/1% BSA were incubated with the samples at room temperature for 1 h. The samples were then washed 3 times in PBS and mounted with Vectashield containing DAPI. Fluorescent images were acquired using a Zeiss Axioskop2 coupled to an MRm (Zeiss) camera at 10x, 20x, and 40x magnifications with Axiovision software. The primary and secondary antibody sources and dilutions were as follows: mouse anti-Nestin from BD Biosciences Pharmingen (1:500), mouse monoclonal anti-Neuronal Class III **β**-Tubulin (TuJ1) from Nordic Biosite (1:500), mouse monoclonal anti-a-smooth muscle actin (SMA) from Sigma (1:1000), rabbit poly-clonal anti-CD133 from Invitrogen (1:500) and rabbit poly-clonal anti-glial fibrillary acidic protein (GFAP) from DAKO (1:500). Species-specific Alexa-488, Alexa-594-conjugated secondary antibodies were used as appropriate and were obtained from Molecular Probes (1:500).

### RT-qPCR

Total RNA was extracted from cells using RNeasy (Qiagen) and contaminating DNA digested using RNase free DNase kit (Qiagen). cDNA was synthesized using 200 ng of total RNA using High Capacity cDNA Reverse Transcription Kit (Applied Biosystems). 1:25 dilution of the cDNA was used for real-time PCR. Platinum SYBR Green qPCR Supermix UDG (Invitrogen) was used for real-time PCR analysis with the 7500 PCR system (Applied Biosystems). Primers (MWG Biotech) are available on request.

### Staining of rat neural stem cells (NSCs)

Neural stem cells were plated in a 6-well plate (40 000 cells/well) in serum-free DMEM:F12 (Invitrogen) with supplements and 10 ng/ml FGF2 (R&D systems) for 24 h prior to further stimulation. Cells were then stimulated with 10 ng/ml FGF2, 10 ng/ml CNTF (R&D systems), 10% FBS (Invitrogen), 1 mM VPA (Sigma) or without added factors (N2 medium) for three days. Addition of soluble factors was carried out every 24 h, and media was changed every 48 h. The LCOs were dissolved in deionized water to a final concentration of 1 mg/ml, and administered at a dilution of 1:500, directly to each well and the detection was done after 10 min in a fluorescent microscope. p-HTMI generated fluorescence at a wavelength common to green fluorescent proteins. A strong green signal was obtained in undifferentiated immature stem cells, accumulated in the cytoplasm of the cells, whereas differentiated and more mature cells displayed a significantly lower or no signal. Neural stem cells were also treated with 10 ng/ml BMP4 and 10 ng/ml Wnt3a (R&D Systems) for 14 days and then stained with p-HTMI.

### Staining of rat C6 glioma

C6 (ATCC CCL-107) glioma is a rat cell line used to as a model system for glioma cells. The cells were grown in DMEM medium (Invitrogen) supplemented with 10% FBS. For the purpose of maintenance, the C6 glioma cells were grown in 75 cm^2^ flasks. Prior to experiments the cells were split and plated (40 000 cells/well) in a 6-well plate. HEK-293 (ATCC CRL-1573), COS-7 (ATCC CRL-1651), and CV-1 (ATCC CCL-70) fibroblast cell lines were cultured according to the supplier’s recommendations.

### Staining of rat C6 glioma cultured as NSCs

C6 glioma cells were cultured with the same protocol as for NSCs, on plates pre-coated with poly-L-ornithine and fibronectin and then grown in N2 medium with supplements. The cells were plated in a 6-well plate (40 000 cells/well) in serum-free DMEM:F12 (Invitrogen) with supplements and 10 ng/ml FGF2 (R&D systems) for 24 h prior to further stimulation. Cells were then stimulated with 10 ng/ml FGF2, 10 ng/ml CNTF (R&D systems), 10% FBS (Invitrogen), 1 mM VPA (Sigma) or without added factors (N2 medium) for three days. Addition of soluble factors was carried out every 24 h, and media was changed every 48 h. p-HTMI was administered, at a dilution of 1:500, directly to each well and the detection was done after 10 min in a fluorescent microscope. p-HTMI generated chemoluminescence at a wavelength common to green fluorescent proteins.

### Two-photon microscopy of NSCs stained with p-HTMI

The emission wavelength of p-HTMI was characterized in a two-photon microscope. NSCs were grown in 35 mm plates (40 000 cells/plate) and were treated with 10 ng/ml FGF2 for 48 hours. Prior to staining the cells, the medium was changed to DMEM:F12 medium without phenol red (Invitrogen) in order to eliminate background signals. CellTracker (Invitrogen) was administered together with p-HTMI (1:500) in order to be able to find living cells, resulting in a double staining in red (CellTracker) and green (p-HTMI).

### FACS-sorting of NSCs stained with p-HTMI

NSCs were grown in 35 mm plates (40 000 cells/plate) and were treated with 10 ng/ml FGF2 for 48 hours. Prior to staining the cells, the medium was changed to DMEM:F12 medium without phenol red (Invitrogen) in order to eliminate background signals. Plates were incubated with p-HTMI (1:500) for 10 minutes and then incubated with HANKs for five minutes. The cells were then scraped and run through a FACS machine. Analysis was carried out on a FACSCalibur flow cytometer equipped with CellQuest software (Becton Dickinson).

### Mouse Intracranial Injections

4 weeks old NOD.CB17-PrkcSCID/J mice (from Charles River) were anesthetized (4% isoflurane) and received a stereotactically guided injection of 250 000 glioblastoma-derived stem-like cells into the right striatum (2 mm lateral and 1 mm anterior to bregma at 2.5 mm depth) in 2 μl PBS. Mice were weighed weekly and monitored for neurological symptoms. Between 5-24 weeks after injection depending on the injected cells, mice received a terminal dose of Avertin followed by a stereotactically guided injection of 4 μl p-HTMI in the same site of the previous surgery for 10 min. Mice were perfused with PBS at first, followed by 0.2% p-HTMI solution in PBS and then with 4% PFA. Brains were removed, fixed in 4% PFA overnight at 4°C and then transferred in 30% sucrose solution overnight at 4°C. After freezing, brains were sectioned (30 μm) using a microtome and stored in anti-freeze solution (30% etylenglycol, 30% glycerol in PBS) at −20°C. Animal experiments were performed in accordance with national and local guidelines (ethical permit no N110/13).

### Immunohistochemistry

Floating sections were washed in PBS, blocked in 10% FBS blocking solution (0.5% glycine, 0.2% Triton X-100 in PBS) for 30-120 min rocking and further incubated overnight with primary antibody solution (1% FBS blocking solution) with 1:1000 mouse anti-human nuclei antibody (MAB1281). On day two, sections were incubated with secondary antibody in 1% FBS blocking solution for 1 h. Sections were then mounted with media containing DAPI.

### Cell culture and FACS-sorting of glioblastoma-derived stem cell-like cells (GCs), U-87MG with p-HTMI, CD133, CD44, CD271, PI, and Annexin V

Patient-derived GBM cells (U3034MG, U3088MG, U3031MG) or GCs were grown in 6-well plates (50 000 cells/well), coated with poly-L-ornithine (15 μg/ml) and laminin (10 ug/ml), grown to a confluency of 80%. Cells were expanded in medium containing Neurobasal, DMEM/F12 glutamax, media supplements and EGF and FGF (10 ug/ml, 1:1000) every 72 hours. The plates were incubated with p-HTMI (1:500) for 10 min, then incubated with Accutase for 7 min and re-suspended in cell culture medium. Cells were spun down at 1500 rpm for 5 min. Further, incubated with binding buffer (BSA, 0.5M EDTA and PBS), (100 μl/ sample), CD133-APC (130-090-826 Mitenyi Biotec) 10 μl/sample, incubated for 10 min at 4°C and washed with PBS prior to FACS analysis. Cells were incubated with CD44 (45-0441 eBioscience) diluted in binding buffer and used at a concentration of 1:10 000. Same antibody conditions as CD133 were used for CD44 i.e. 100 μl binding buffer, incubated for 10 min at 4°C and washed with PBS prior to FACS analysis. Cells were also double stained for p-HTMI and CD271 (560834 BD Pharmingen), 5 μl/sample with the same antibody conditions as CD133. U-87MG cells were expanded as previously described^30^ and incubated with the same conditions of p-HTMI and CD133, CD44 as described for GCs. To verify cell necrosis, lysis buffer was added with 5 μl PI per sample and for cell apoptosis, cells were incubated with 100 μl 1X Annexin V binding buffer, 5 μl APC Annexin V per sample (561012 BD Pharmingen) and incubated for 15 min. The analysis was carried out on a FACS LSRII flow cytometer equipped with FACSDiva software. Patient tissue collection and use were in accordance with ethical permit EPN Uppsala 2007/353 and its addendum Oct. 28, 2013.

## Author Contributions

S.I., A.G., and B.M. pursued the stem cell and glioma stem cell experiments, including cell culture, immunocytochemistry, immunohistochemistry, FACS, microscopy, RT-qPCR etc. A.K.O.Å, M.B., R.S., P.K., and K.P.R. generated and produced the LCOs including p-HTMI. E.K. and B.J. provided infrastructure and assisted in FACS sorting. S.N., B.W., L.U., and K.F.N. provided human glioblastoma cells. A.I.T. pursued the initial screen of LCOs in neural stem cells together with S.I.. V.R. and J.H. performed the *in vivo* generation of tumors in mice. P.U. supervised the two-photon microscopy. K.P.R and O.H. designed and directed the study, analyzed the results with the colleagues, and wrote the manuscript.

## Figure legends

**Scheme 1.**
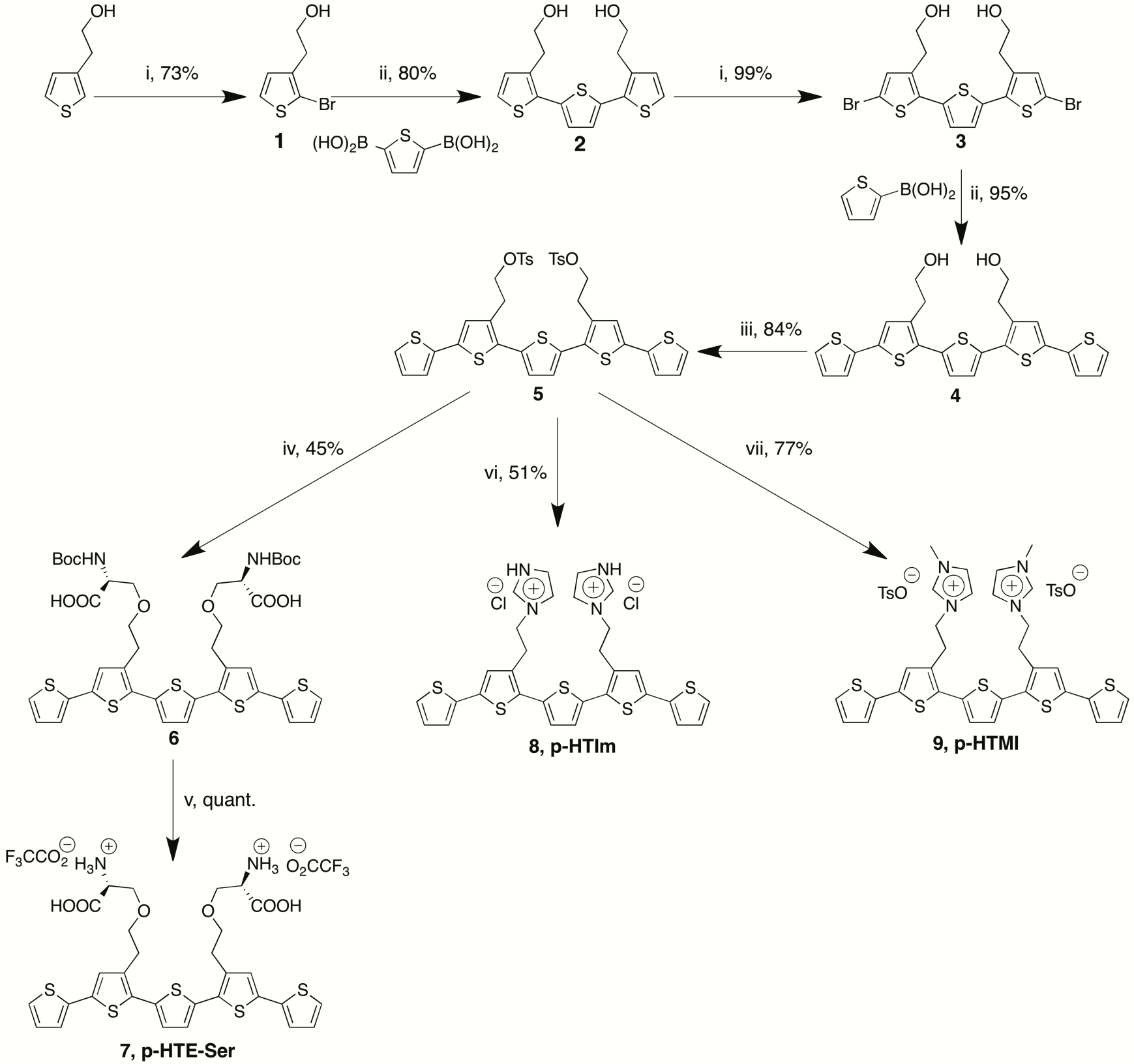
Synthesis of p-HTMI. Reagents and conditions: (i) *N*-bromosuccinimide, DMF, −15°C; (ii) PEPPSI-IPr, K_2_CO_3_, toluene/methanol (1:1), 75°C; (iii) *p*-toluenesulphonyl chloride, CHCl_3_, pyridine; (iv) methylimidazole, acetonitrile, 75°C; (v) imidazole, acetonitrile; (vi) Boc-L-Ser-OH, K_2_CO_3_, DMF, 50°C; (vii) TFA, CH_2_Cl_2_.

## Supplementary Figure legends

**Supplementary Figure S1:**
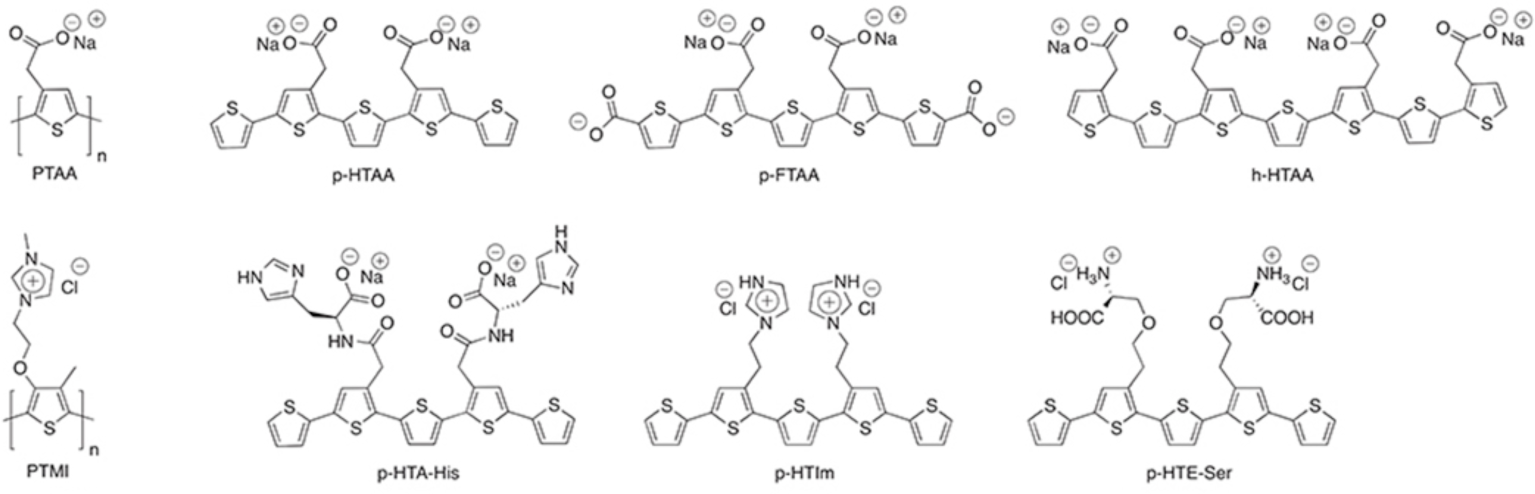
Structures of the LCOs evaluated.

**Supplementary Figure S2:**
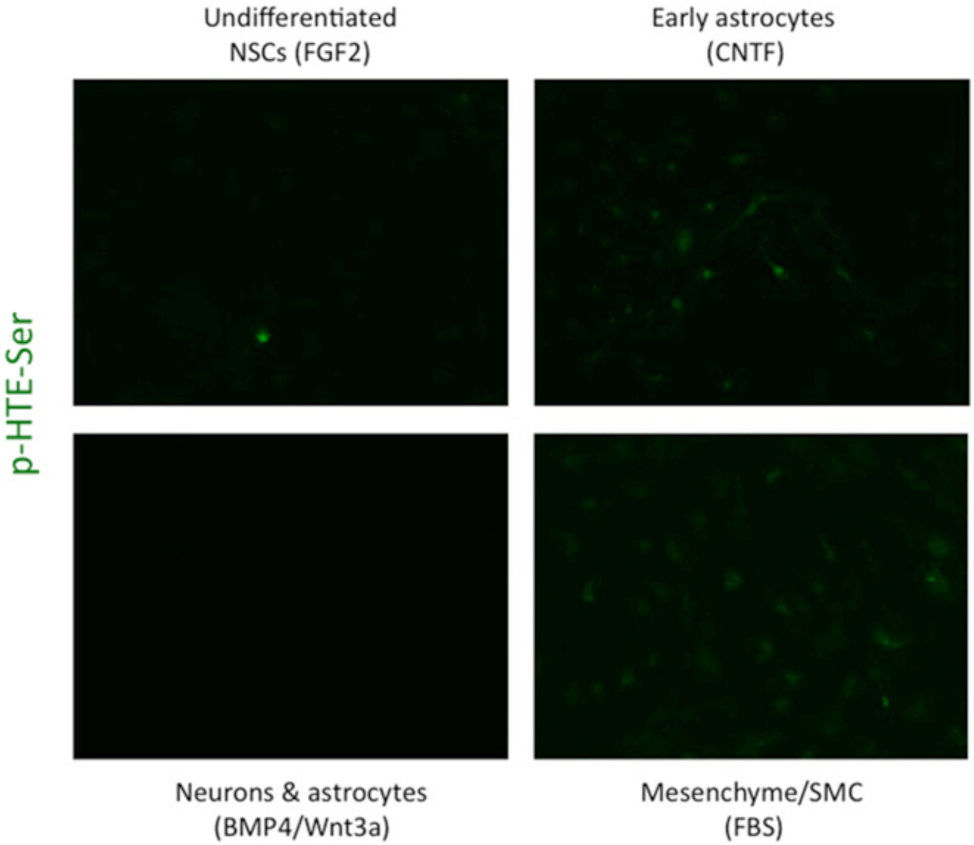
A different LCO, p-HTE-Ser, does not stain NSCs, while a faint labeling can be detected in NSC-derived early astrocytes and mesenchyme/smooth muscle like cells.

**Supplementary Figure S3.**
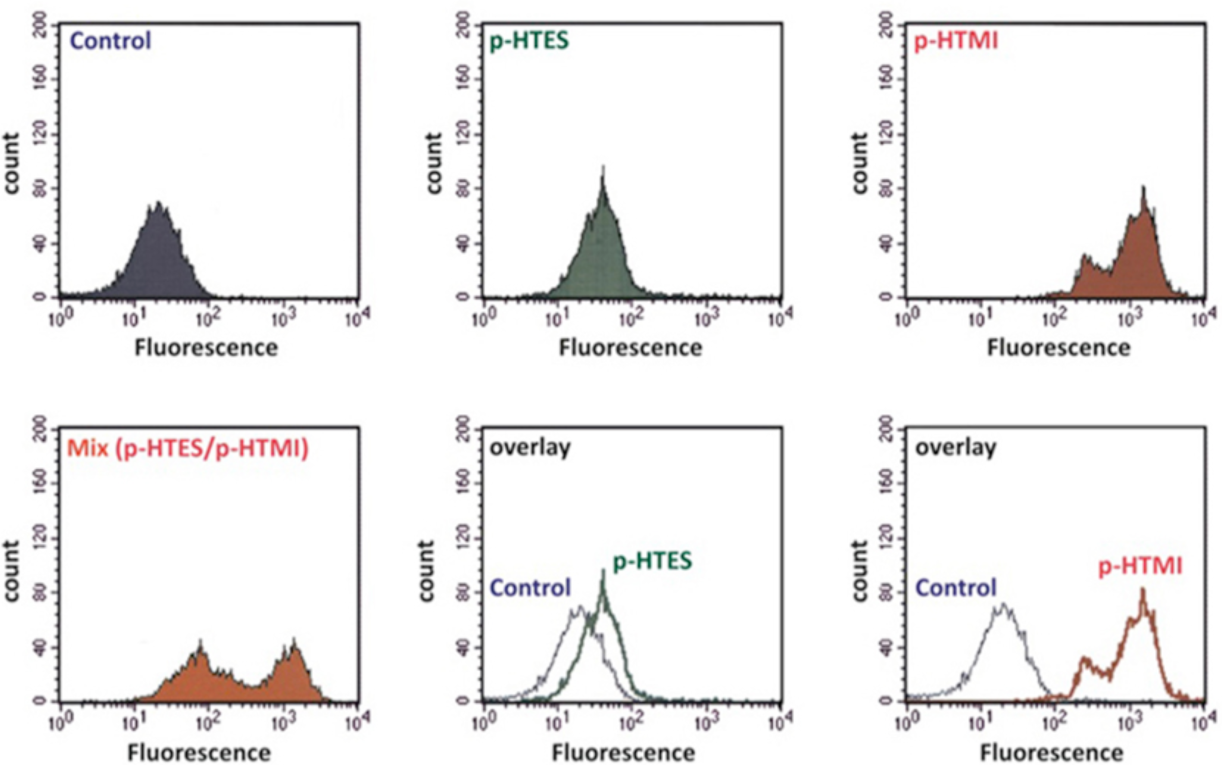
p-HTMI, but not p-HTE-Ser, efficiently and selectively labels NSCs in mixed cell populations as assessed by FACS. p-HTE-Ser shows a similar distribution as control cells which has not been incubated with an LCO, whereas p-HTMI specifically detects the NSCs, also in a mixed population of cells (log scale).

**Supplementary Figure S4:**
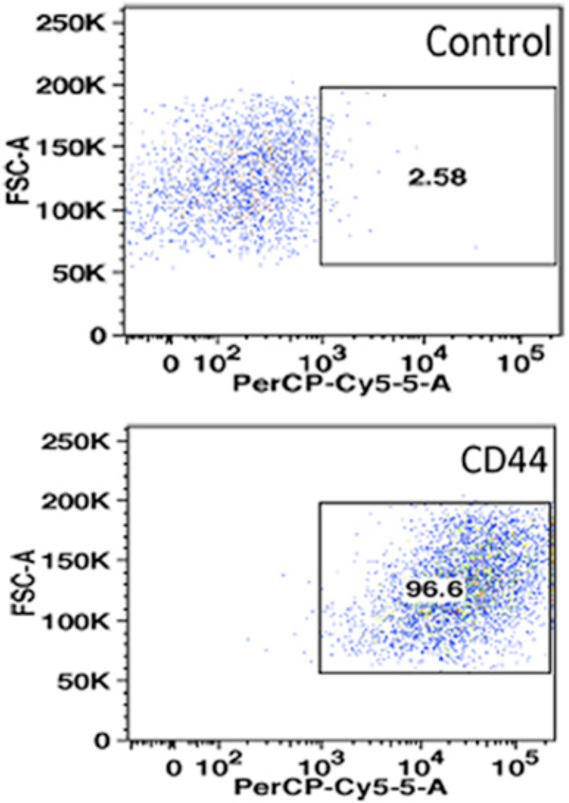
FACS analysis suggesting low specificity of CD44 in GCs

**Supplementary Movie M1:** Stack of micrographs from two-photon microscopy demonstrating that the p-HTMI labeling was predominantly cytoplasmic in NSCs *in vitro*. Blue=DAPI, green=p-HTMI.

**Supplementary Movie M2:** Stack of micrographs from confocal microscopy demonstrating that the p-HTMI labeling was predominantly cytoplasmic in human tumors in mouse brains *in vivo.* Blue=DAPI, red=HuNu, green=p-HTMI.

